# Envelope Simulation: A New Adaptive Method for Dose Optimization

**DOI:** 10.1101/2020.03.09.983411

**Authors:** Steven Piantadosi, Guohai Zhou

## Abstract

We present a flexible and general adaptive experimental design for dose finding clinical trials. This method can be applied in sterotypical settings such as determining a maximum tolerated dose, when the dose response relationship is complex, or when the response is an arbitrary quantitative measure. Our design generalizes dose finding methods such as the continual reassessment method (CRM), modified CRM, and estimation with overdose control (EWOC). Similar to those, our design requires a working mathematical model, which assures efficiency, low bias, and a higher fraction of dose selected near the optimum compared to purely operational designs. Unlike typical model based designs that always employ the same model, our method allows individual dose response models tailored to the circumstance. Simulations are also integral to our design allowing it to account for both known and unknown effects of dose on outcome. This method is applicable to general dose finding problems such as those encountered with modern targeted anti-cancer agents, immunotherapies, and titrations against biomarker outcome measures. The method can also support improvements in the design of the experiment being conducted by providing a platform for concurrent simulations to assess the influence of projected design points. Although we present the design in the context of clinical trials, it is equally applicable to experiments with non-human subjects.

## Introduction

This paper introduces a general algorithm for design and execution of dose finding experiments. Many study designs for the restricted task of determining the maximum tolerated dose of a cytotoxic agent in oncology have been proposed and implemented in the literature. However, methods general and flexible enough for dose optimization in the broadest context are lacking. Because of the diversity of such questions it is not clear that a universal method exists. However, a combination of flexible techniques as discussed in this paper allows considerable unification toward a general method for dose response optimizations.

Useful lessons regarding dose optimization are evident from the narrow and well studied problem of dose escalation for oncologic therapeutics where a “more drug is better” philosophy has been the rule. The designs used for that purpose fall into two classes: empirical (or operational) and model-guided. Empirical designs are typified by the “up-and-down” or “3+3” dose escalation, named for the cohort sizes frequently employed at each dose level. These designs have their origins in ordinance fuse testing during World War II and have had minimal modification in the last 75 years. Small improvements in performance have resulted from optimal spacing of prespecified doses (the so-called Fibonacci escalation), and employing single subject cohorts early in the escalation (accelerated titration). Unfortunately, these designs have poor operating characteristics when applied to true dose finding problems, such as locating the dose associated with a prespecified frequency of toxicity (maximum tolerated dose or MTD).

Model guided dose escalations were introduced 30 years ago [1] and are typified by the continual reassessment method (CRM) and its relatives. Both the modified CRM and estimation with overdose control (EWOC) [2] are in this class of designs, because among other things they both use dose-response models to interpolate and extrapolate data to estimate the target dose. Model guided designs are more efficient and have superior operating characteristics compared to empirical designs [3, 7]. In this paper, we will not review the important nuances of these historic designs. Our focus is on broadly applicable methods.

Empirical designs remain widely used in cancer therapeutics development and are often endorsed implicitly or explicitly by important research sponsors such as FDA and NIH. However, model guided dose escalation has been shown to require fewer subjects, have no bias towards low doses, treat a higher proportion of study participants near optimal doses, and outperform empirical methods in essentially all performance domains that have been tested by statisticians [7]. The simplicity and clinical independence offered by empirical designs appears to overcome operational superiority in the minds of many. Outside cancer we do not routinely employ treatments with a high frequency of side effects. An optimal biologic dose for antibiotics might be a maximum nontoxic dose. For antihypertensives and analgesics, the optimum might be a minimum effective dose. A good example is aspirin: when used as an anti-inflammatory the dose needed could be as high as maximum tolerated if the underlying severity of the condition requires it. As an analgesic, aspirin dose is maximum nontoxic. As a preventive, the dose is minimum effective.

Inferior and inappropriate designs are often applied where they can’t succeed. This can happen when empirical dose escalation designs are uncritically applied to more general questions, especially outside the domain of cytotoxic drugs. But the absence of general dose finding methods pushes clinical investigators to employ standard designs even when they are not suitable for the underlying question. One limitation of MTD study designs is that the response scale is a probability. However, many essential questions require quantitative responses that are not probabilities. Examples include immunological reactivity, pharmacokinetic outcomes, target binding, or tissue concentrations. Translational needs such as optimizing dose for quantitative biomarker outcomes is another example. There have not been any experiment design methods that facilitate optimal dose finding in such circumstances.

## Methods

### Requirements

A study design widely suitable for dose finding would have to perform well for diverse dose optimization questions such as a) classic dose escalation to locate the MTD, b) location of the dose associated with a peak response, c) location of a dose that nearly saturates a response, d) titration to a prespecified measured outcome under arbitrary dose response relationships, and e) location of the lowest dose that yields a prespecified response level. A general dose finding method should also incorporate relevant background information, opinion, or intuition from the clinical investigator. It must make efficient use of data and be ethically appropriate for study participants. Also it must have simple, transparent, and flexible operational rules. Finally, a general dose finding method should have reliable, simple, and robust software to support it.

The method we present satisfies these requirements. Our method contains both deterministic and stochastic components. The deterministic portion is model guided, but is not restricted to a single model. Models confer efficiency and accuracy under fairly general conditions, but also allow flexibility because the mathematical construct can be tailored to essential features of the problem. A monotonically increasing model might be appropriate for MTD questions, however one with a peak and fall-off would be more appropriate for questions about dose maxima. Any model that mimics biological behavior without being restrictive could be useful in appropriate places.

Our method is agnostic with regard to the actual model used. To augment efficiency and flexibility, the stochastic part of our method incorporates statistical simulations to characterize variation. These intrinsic simulations represent a range of stochastic behavior that investigators anticipate, but are difficult to capture quantitatively. For the deterministic component, the investigator selects an appropriate dose response model. For the stochastic component, the investigator specifies regions or envelopes in the outcome space from which actual data could arise. Data sampled from the envelopes substitute for prior specification of model parameters as in a Bayesian method.

To initiate dose finding, data are sampled randomly from the given envelope regions and the model is fitted; repeating this simulation process a large number of times yields a distribution of fitted features. The distribution captures both elements of the problem specification, and allows the investigator to choose a starting dose in light of both deterministic and random behavior. Updating this simulation process with actual data as they are collected creates a learning algorithm conditional on the design assumptions. We refer to this as Envelope Simulation (EnSim) guided dose finding.

### Algorithm

The basic EnSim algorithm is shown in Table 1. Steps 1-4 are preparatory. Steps 5-9 iteratively update results with new data. During this process, the envelope data diminish in influence in favor of the real data being collected. Step 10 is optional but offers the opportunity to strengthen the design of the trial. Details regarding each step of the algorithm are provided in the following sections.

**Table 1.**
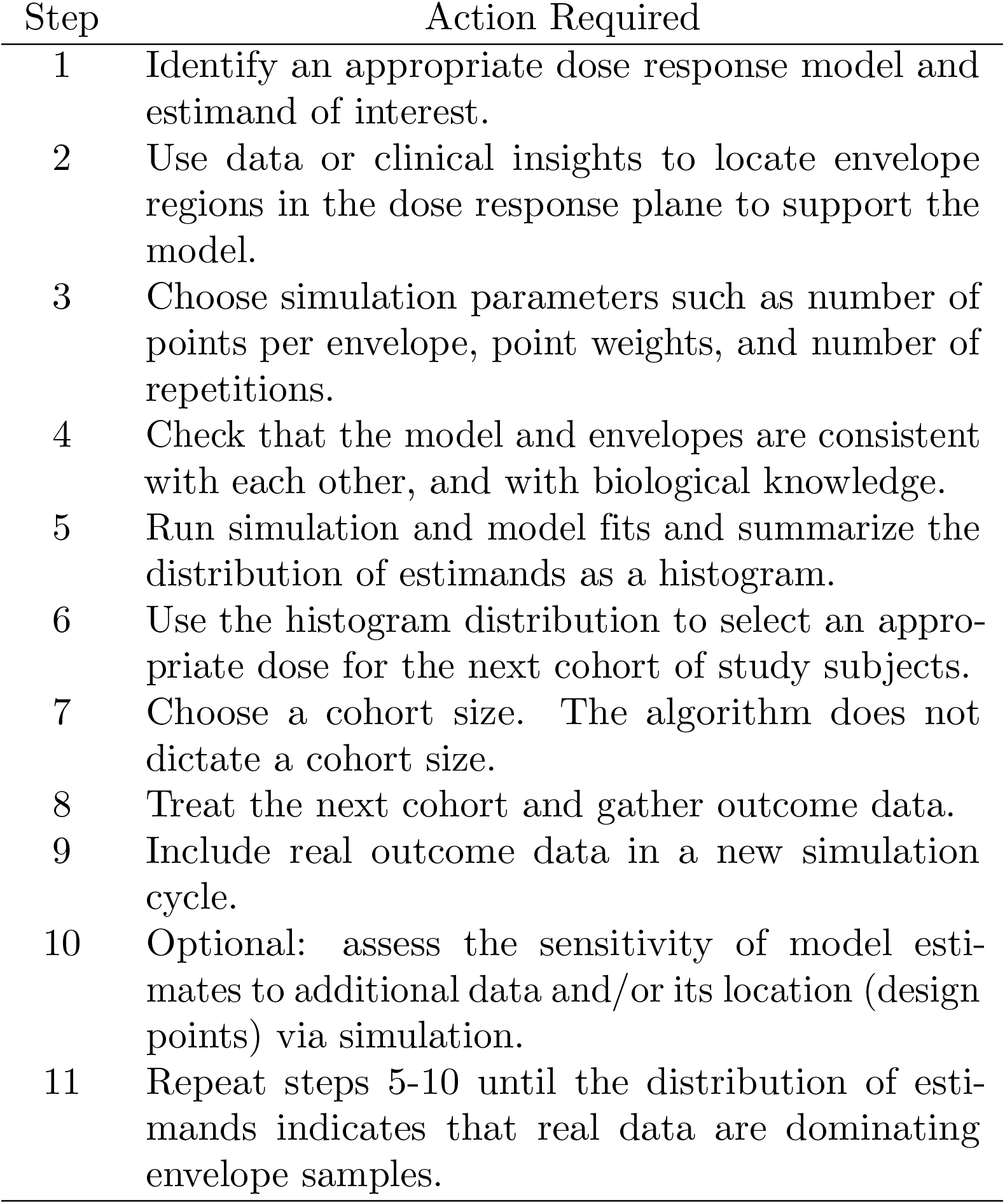
Steps in the Envelope Simulation Algorithm.

The flexibility of the algorithm is its main advantage. Investigators must choose wisely with regard to the biological model, envelopes, simulation parameters, subject cohort sizes, and dose. The list is not very different from the requirements and subjectivity surrounding operational designs. Familiar operatonal designs can appear less subjective but only because there are traditional choices associated with them. Traditional does not imply optimal.

### Specifying a Model

We have tested a number of dose response models in the EnSim algorithm. For all models, dose is represented by *d*. Any model that guides dose finding should represent biological knowledge of the problem without being unnecessarily restrictive. No single dose response model can suffice for all purposes, but an algorithm for implementing a model can be nearly universal.

Because of its sigmoidal shape, the logistic equation is a classic dose response function:

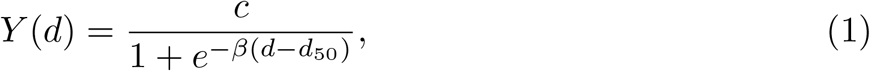

where *c*, *β*, and *d*_50_ are parameters to be estimated from data. The parameter *c* represents the maximal response. If we restrict *c* = 1, *Y* (*d*) can then be taken as the probability of response. The logistic model has often been used in that form for determining the MTD. Dose escalation can be initiated by specifying probability distributions for the parameters (Bayesian method), or using pseudo-data [4] that serve the same purpose. Dose escalations tend to be more about location (*d*_50_) than slope (*β*), and many alternative models might be employed for MTD determination, including a simple line segment. The model does not have to be “true” across its full range – only approximately correct locally for such methods to work nicely.

The logistic and similar models are monotonic and cannot represent true peak responses. This biologic behavior could be seen with immunologic agents for example. Features of such a shape could include a rising slope, an inflection point, a peak, a post-peak plateau, and a dose scale factor. Hence we might anticipate that a model for this purpose could incorporate five or more parameters. A flexible model that contains this behavior is

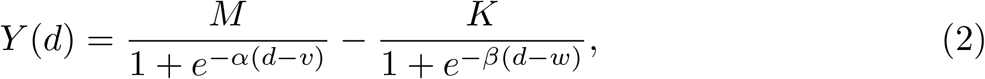

where *M*, *K*, *α*, *v*, *β*, *w* are free parameters with *M > K*. One parameter might be eliminated by setting *α* = *β* without losing much flexibility. The response *Y* (*d*) is not required to be a probability in such a model. A five- or six-parameter model is capable of a wide range of behavior and would have to be supported by substantial data both during its initiation and when drawing biological conclusions. The use of this model will be illustrated below.

If the biological question requires finding the maximum dose response, one might consider a simple model with a well defined maximum, even if it is not derived from underlying biology. One such model is a parabola, which points in the correct direction if parameterized as

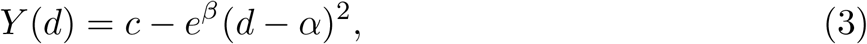

where the peak will be at *α*, and *e*^*β*^ is guaranteed to be positive. This model has only 3 parameters. Finding a peak using a model is not purely a location problem because of the shape of data on each side of the peak, so the stiffness of a parabolic form may be a problem.

Other mathematical forms that can point to a response maximum at some dose *α* are bilinear models with a breakpoint at the maximum. These include

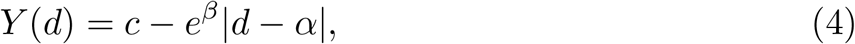

which is symmetric, and

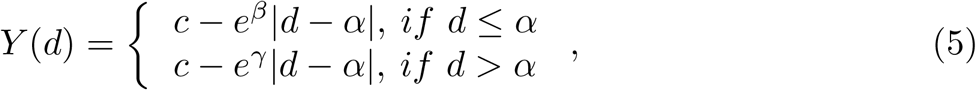

which is asymmetric around *α* by virtue of a fourth parameter. This last model can be an efficient locator of maxima despite its lack of biological realism, as illustrated below.

A model that strictly saturates a maximal response *c* is

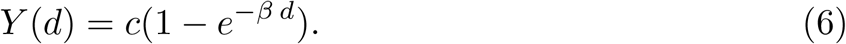

With *c* = 1, this model can also represent a probability of response. Because *Y* (0) = 0, it does not represent a background or a threshold.

The allometric form

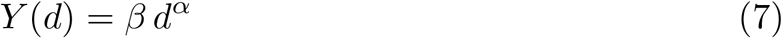

is another simple, flexible, biologically grounded descriptor of general relationships.

A final flexible model that embodies useful behavior is

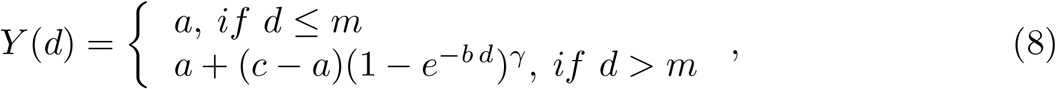

where *a* represents background response, *m* is threshold dose, and *c* is the maximal response. This is quite a flexible model that comes at the expense of five parameters.

The message here is not that any of these is a universal model, but that the EnSim algorithm can incorporate any deterministic part provided that the model a) is tractable when fitting data, b) is not over-parameterized, and c) can be supported by data from the experiment. In other words, we have to be able to fit our chosen model to relatively sparse data. The purpose of dose finding is not model validation, but is to extract biologically useful features from data using tools that facilitate without interfering. The model eventually takes a background role to the data. Key inferences are not likely to depend on nuances in the model.

Parameterization is always a concern when model fitting. For the models above, certain parameters must be positive for the equation to make biological sense. For example, the scaling factor for dose should be positive the way the models are written. The asymptotic response at high doses should also be positive for these models, although this does not have to be true universally. In Eq 2, it should be the case that *M* > *K*. When model fitting, these types of constraints can be added explicitly as ancillary equations. Alternatively the models can be parameterized to force constraints implicitly. For example, replacing parameter *b* by *e*^*β*^ assures that a positive coefficient always results from model fitting. The computer program allows either explicit or implicit constraints, although the latter appear computationally faster.

### Specifying Envelopes

The stochastic portion of the EnSim dose finding method relies on user-specified envelopes, which are a new concept. Envelopes are regions of the dose-response plane from which we expect data to arise. The expectation is founded on clinical or biological knowledge and experience prior to data being gathered. Consider this hypothetical circumstance: if a new drug is administered at a dose between 100-200 units, the response will be 10-20 measured units; if it is administered at doses between 300-400 units, the response is expected to be 30-40 units. Such expectations might come from knowledge of similar agents, preclinical data, or clinical data at other doses or in other cohorts. The implied envelopes in the dose-response plane have corners at {(100, 10), (100, 20), (200, 10), (200, 20)} and {(300, 30), (300, 40), (400, 30), (400, 40)}. We might expect the model to pass through these regions absent other information or data (Fig).

How do investigators construct a set of envelopes for a given problem? This question is reasonable in the context of a new design. Interestingly, elicitation of key design components is not typically addressed when using traditional designs. For example, in a 3+3 design, where do the proposed doses come from? In a fully Bayesian CRM or EWOC design, where do the prior probability distributions for model parameters come from? We seldom ask how IRBs, Ethics Boards, sponsors, or regulators know that a given set of doses is appropriate for the study population. Nor do we usually ask how reasonable are the hypothetical alternatives used to describe operating characteristics for a given design. There is no extensive literature dealing with such questions – the answers are judgment based. When investigators have sufficient knowledge and experience with the clinical context, the design space/behavior, and the class of agents under investigation, it is not difficult to approximate the dose response relationship. Nor is it troublesome to hypothesize an appropriate mathematical model. Reasonable starting conditions are all that is necessary because real data will quickly dominate in the algorithm.

Envelopes embody both knowledge and ignorance. They do not literally specify dose-response pairs – that would seem to indicate strong knowledge. Sketching a region, all points of which are equally likely to contribute data, characterizes also what we do not know. Low weights or utilities can be assigned to simulated data from an envelope so their influence on model fitting is small compared to real data. This further emphasizes our ignorance about the truth of nature. Although the influence of envelopes will ultimately be small in the presence of real data, they are essential 1) early in the dose finding algorithm to specify the model, and 2) to support the model in regions where actual data are sparse or cannot be obtained.

Several envelopes need to be specified at the initiation of a dose finding trial. Their locations must be consistent with the shape of the chosen model, and constrain it to plausible regions. In some cases, envelopes might overlap. But care is needed because only a few hypothetical observations will be drawn from each envelope. We do not want to create inconsistencies between the simulated data and the ability of the model to represent them. Fig shows a set of envelopes and a model that is consistent with them using Eq 2.

There is no requirement that envelopes be rectangular, although all the examples in this paper make use of them. Rectangles are simple and offer a small advantage in simulation efficiency compared to regions of other shapes. This implies that the dose-outcome random sample will have a bivariate uniform distribution within each envelope. Other region shapes or different bivariate distributions could be used, but doing so increases complexity without promise of improved performance. One of our motivations is to avoid assumptions regarding prior distributions for multiple model parameters as would be required in a formal Bayesian method. Hence complex envelope sampling distributions are unwelcome.

The initiation of the algorithm relies on sparse samples from each envelope. Fitting a multiparameter model to sparse data is delicate, and model fits might be poor or impossible in some simulation replicates. This problem can be reduced or eliminated by sampling more points from each envelope or from the most influential ones, adding more envelopes, making them smaller, or making them jointly more consistent with model behavior. The number of envelopes should be the same or greater than the number of model parameters, and they should be placed widely. A small fraction of failed model fits will not damage the overall process because real data ultimately dominate. The algorithm can succeed in the presence of a large fraction of failed fits, but it may then be inefficient. Such a circumstance indicates structural problems with the experiment plan, envelopes, or model, and is simple to avoid.

Each clinical problem requires a unique set of envelopes. Different investigators might choose different envelopes for the same problem. However, in the course of conducting the trial, data from the subjects treated will control the model fit, and overcome low-weighted samples from the envelopes. When there are at least as many actual data points as model parameters, investigators could remove the envelopes and rely on real observations for subsequent model fits. Even at that point, a mix of actual and simulated data may be needed because the experiment may not gather real data from certain dose ranges. Simulated data from those ranges will not be strongly influential in the region of interest, but may be necessary for a good overall model fit. Removing envelopes converts the algorithm essentially into a generalized CRM.

We do not illustrate it in this paper, but it would be simple to weight envelope data dynamically as the number of real data points increases. For example, the weight of envelope data could be taken to be 1/(1 + *n*) where *n* is the number of real data points. As with a fixed fractional weight, this guarantees that final model fits will be dominated by real data.

As the influence of envelopes vanishes we will be left with ordinary model fitting issues. These include extracting the estimand or feature of interest, precision in estimating parameters, lack of fit, and bias due to lack of data in certain ranges. Resource limited dose finding studies yield sparse data unlikely to overcome all these problems easily. Simulation mitigates the difficulties. The limitations of sparse data are universal and should not be taken as shortcomings of any underlying method. Adequate real data will overcome a multitude of problems, and could even indicate deficiencies in the model being employed. It is perfectly reasonable to choose a better model at any time the data indicate, but those cases will not be illustrated in this paper.

## Results

The deterministic and stochastic parts of the EnSim algorithm are combined in simulations. In a simulation, observations are randomly sampled from each envelope, combined with available real data, and the model is fitted via non-linear least squares or a similar method. Simulated data are assigned low weight in model fitting.

### Estimands

The estimand is the data or model feature estimated following the model fit. For a classic cytotoxic dose escalation design, the estimand might be the dose at the intended quantile, for example the dose that yields a 30% probability of a dose limiting toxicity. For detecting a maximum response, the estimand would be the dose under the peak of the fitted model. For measured biomarkers, the estimand might be the dose that yields a prespecified level of response.

The estimand might be obtained using any of three techniques. All three methods employ the parameter estimates from the current simulation replicate as though they were fixed constants. One method is algebraic calculation from the model parameter estimates. Inversion of the logistic model to yield dose as a function of quantile response is an example of this. This works only when the model has a closed form inverse. Another technique is a secondary numerical calculation from the fitted model. Finding a maximum or minimum is an example. This approach is based on root finding numerical algorithms, and is general enough to cover the first case as well. Lastly, the estimand might be available directly in the form of a model parameter. The bilinear models for maxima discussed above (Eq 4 and 5) are examples where the maximum is represented explicitly by the parameter *α*.

Simulation replicates yield a distribution of estimands that represents sampling variation from the envelopes combined with uncertainty in the model parameter estimates. Investigators can examine that distribution and choose the next dose or design point accordingly. The chosen point need not be a particular quantile of the simulation distribution, such as the median. It should be chosen simply to be consistent with the current data and the specifications of the deterministic and stochastic parts of the experiment setup. This will be illustrated below. This process is repeated with real data from each new design point, and converges quickly under very general conditions. Convergence can be seen by the narrowing of the sampling distribution when actual data dominate the model fit, as illustrated in the next section.

Early termination of the algorithm is possible. Such a method ideally would be based on the value of information to be gained by continuing. Developing such methods is outside the scope of this paper.

### Simulation performance

All of the models described above have been used in simulations of the algorithm, and yield similar qualitative results. Here we describe two examples that illustrate key findings. First is a dose finding example using the logistic model from Eq 1. The following section covers a second example where the problem is locating the dose associated with a maximum response, and employs a model without a biological rationale.

The process of conducting the algorithm is shown in Figures and for a three-parameter logistic model dose titration to a measured response value of 4.0. The dose and response units of measurement are immaterial to the algorithm. Note that the response scale extends from 0 to 10, indicating a measured response that is not a probability. In this example, 1000 simulations were performed at each step with samples of size 3 from each envelope. The ordinates of the estimand histograms are uniform, but the abscissas are not. The histograms narrow dramatically when the actual data can support three model parameters. This indicates the declining influence of the envelopes in the presence of real data. At the final model fit in Fig, the estimated model parameters from Eq 1 are *c* = 9.11 ± 1.12, *β* = 0.07 ± 1.25 and *d*_50_ = 52.4 ± 1.08. Five data points appear to determine the model fit with excellent accuracy.

Although the variability attributable to sampling envelopes vanishes, uncertainty always remains because of intrinsic (subject to subject) variation. That variability is revealed in the fitted model parameters. Even when real data dominate the simulated data, the model can still fit poorly, or the data may be barely able to support the number of parameters in the model, or resolve nuances in its shape. One could not claim in such cases that being free from envelope simulated data is a complete solution to the problem. More data will always help unless the model is systematically inappropriate.

Some issues surrounding the design remain. These include the cohort size, number of sample points to draw randomly from each envelope, whether or not they should be the same for each envelope, how to assign weights for simulated data in model fits, and how many simulations to perform. In developing and testing this methodology, some answers to these questions have emerged. Because each real observation has much higher weight than an envelope sample, relatively small cohorts from 1-3 observations work nicely. If for some reason envelopes are given more weight at initiation, cohort size should probably be increased to aim for a ten-fold or higher ratio. Assume there is one appropriately located envelope for each model parameter. To initiate dose finding, between 1 and 5 sample points from each envelope should be sufficient. This might seem like a lot of data, except that the weights are low. Absent real data, the weights are irrelevant, and the initiating model fits will work well.

Some envelopes may have more leverage than others in a model fit. If this is an issue, or if real data are gathered in the close region of an envelope, the number of points or weight of that envelope data can be set to zero subsequently. This flexibility is inherent in the algorithm. A workable guideline is that simulated data should have about 1/10th the weight of real data. Weights can be specified by the user in the computer algorithm and need not be constant over all iterations. A number of simulations between 100-1000 seems to work well. This provides a reasonable resolution of the sampling distribution without consuming much computer time.

With sparse data there usually is insufficient information to indicate mis-specification of the model or envelopes. In dose escalations, mis-specification can be seen if the high-dose high-response envelopes are set too low. Real data might then readily show inconsistency with those envelopes in the “upper right” region of the dose response plane. Practical considerations suggest that the best course of action is simply to specify or revise appropriate envelopes given the information at hand, even if some of that information comes after the start of the experiment. Detecting model mis-specification formally requires substantial data. However, we know from the satrt that our model is not literally true. Because the model only needs to be locally approximately correct, and it takes large amounts of information to reveal mis-specification, this is not likely to be a problem. Just as for envelopes, it makes practical sense to replace the model whenever the data suggest that a different one performs better. There is no memory in the algorithm for either envelopes or models any more than there is for previous parameter values.

## Operational Behavior, Software and Extensions

The theory, principles, and algorithm outlined above have been assembled into a computer program to perform all needed calculations and simulations. The program is written in Mathematica [5] and is available with this publication. Mathematica is an interpreted high level language with extraordinary numerical, analytic, and graphics capabilities. The calculations here are nearly trivial for this package, but its power greatly shortens development time. The structure of the language makes insertion of a new deterministic model quite simple compared to a compiled language. Most models are a single line of code, and parameter estimation is accomplished with a single call to the built-in function for nonlinear model fitting.

A computer program written in C++ for Windows is also available for beta testing from the author. This program has a small suite of models (n=12) prespecified. Envelopes and subject data enter the program though a simple GUI. This program is straightforward and fast, but cannot be modified by the user. Because of its speed, it might be particularly useful for investigators needing simulations to optimize experimental designs. Source code is not presently available. We have also implemented the EnSIm algorithm in R. Example 2 below will illustrate it. The R code is available with this publication.

### Example 1

Fig illustrates a hypothetical trial to locate the dose associated with a maximum response. The model chosen is Eq 5, which illustrates how an imperfect model can nevertheless yield the feature of interest. This model has 4 parameters that represent peak response (*c*), dose at peak response (*α*), slope for dose response below peak (*e*^*β*^), and slope for dose response above peak (*e*^*γ*^). The exponential forms for *β* and *γ* are implicit constraints to keep those coefficients positive, guaranteeing that the model will yield an inverted V shape. The top panel in Fig shows the envelopes chosen and initial model fit. Two samples from each envelope with weights 0.1 were used. The data points were each given weight 1. The resulting histogram of estimated peak doses is broad, and even includes a few outliers not shown.

Reliable estimation of the critical dose requires data on both sides of the peak. We assume that several design points to the left of the peak were selected and measured. While an actual trial such as this might carry ethics implications for the order and level of doses, we ignore them here to focus on the methods and software. After fitting data gathered to the left of peak, the results are shown in Fig, middle panel. Intermediate steps are not illustrated. The resulting histogram of estimands has narrowed, but remains imprecise.

Similarly, cohorts were then treated at doses to the right of the apparent peak. The final results are shown in Fig, bottom panel. The histogram of peak dose estimates becomes very sharp, indicating that no influence from the envelope samples remains. For this final model fit, the numerical parameter values and standard errors are shown in Table 2. Because of the way this model is parameterized, the estimand is the parameter *α*. We can see there remains some imprecision associated with the estimate of *α* due to the relatively small size of the experiment.

**Table 2.** Final bilinear model parameter estimates (Eq 5) for the example in Fig. Data from envelopes have been excluded from this fit.

| Parameter | Estimate | Std Err |
| --- | --- | --- |
| $\beta$ | -3.46 | 0.14 |
| $\alpha$ | 48.3 | 3.51 |
| $c$ | 1.51 | 0.07 |
| $d$ | -4.14 | 0.13 |

### Example 2

This example applies EnSim to an actual clinical trial from the literature. The trial in question is a dose escalation study of O^6^-benzylguanine (O6BG) for patients undergoing surgery for malignant glioma [9]. The original data were used as an example by Ivanova and Kim [10] and we use data from their Table 1. The doses of O6BG are in mg/m^2^. Responses are measured alkyltransferase tumor levels in fmol/mg protein. The goal is to estimate the dose level that yields a response of 5 fmol/mg protein as in Ivanova and Kim [10]. The trial starts with a dose of 40 mg/m^2^ and the dose can be escalated in increments of 20 mg/m^2^. Patients are treated in cohorts of three. Given the expected reduction in tissue levels with increasing dose we have selected the model

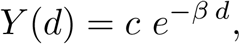

where *d* represents dose, *c* is the concentration at zero dose, and *β* is a decay constant. This equation is similar to Eq 6 above but represents a strictly decreasing function. This new model further illustrates the applicability of the algorithm to any quantitative relationship between dose and response. Furthermore, R demonstrates the use of the algorithm in a widely used statistical tool.

An aggressive design would move to the predicted best dose after each cohort treated. Inverting the model equation yields an estimate of that dose,

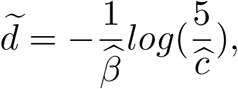

where 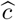 and 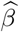 are the current parameter estimates from model fitting to the data. However, the investigators chose a conservative strategy of incrementing doses in a pre-planned escalation. We will also conducted stepwise fits to the cohort data after initiating the algorithm with two envelopes, one at low dose and a second at high dose. Our EnSim algorithm used 1000 simulation iterations, and 3 simulated data points per envelope each with weight 0.1.

Results in Fig illustrates that with increasing data, the model suggests that the optimal dose should be 100 mg/m^2^ with increasing confidence. The actual trial follows a common overdose control policy, i.e., no skipping of untested dose levels. Therefore, the actual dose allocated to the second and third cohorts respectively are 60 and 80 mg/m^2^ instead of the model suggested 100 mg/m^2^. After incorporating the fourth cohort’s data, the EnSim algorithm asserts the theoretical optimal dose for the target response of 5 fmol/mg protein to be 102 mg/m^2^ (95% CI 101-103), and the response corresponding to the optimal administrable dose 100 mg/m^2^ to be 5.3 fmol/mg (95% CI 5.1-5.5). This quantitative summary of results cannot be conveniently obtained by previous methods. Nor is it available with data from eight additional patients and the response-dichotomization method in Friedman et al. [9] or with additional data from eighteen patients.

### Design Optimization

EnSim can assist with design optimization. In example 1, we might ask how to resolve remaining uncertainty regarding the dose at peak. Should a new cohort be treated at the apparent peak, or at some other dose? What is the best size for a new cohort? Additional calculations or simulations can help answer such questions and refine the overall experiment design.

Two qualitatively different approaches are possible. One is to assess the impact of existing design points on either the estimand or summary statistics for the model fit. New points could then be placed in regions of high impact. A second approach is to postulate new design points and assess the likely impact of each point. Response data for a new design point would have to be simulated either from the model or from a new envelope constructed for that purpose. Both approaches will be briefly illustrated below.

Optimizing the design in these or other similar ways can be done at any point there is a stable model fit. This could begin early in the experiment when the model fit depends on envelope data, or later when the fit is determined almost exclusively by real data. The overall method is flexible and adaptable to the circumstances at hand. For example, in a large laboratory experiment, we might try optimizing the design points early to cut down on size and cost. In a human trial, we might apply attempts at optimization late in the design when there is more certainty regarding doses and their effects.

#### Influence of existing design points

With regard to the data in Fig, we can calculate the influence of each existing design point on 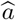, the overall estimate of *a*. This is accomplished by removing a single design point and calculating the resulting estimate, denoted by 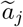. The influence of the *j*^*th*^ design point is defined as

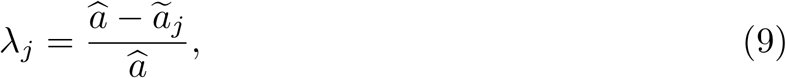

which is the fractional change or sensitivity when the respective point is omitted. It might make sense to gather additional real data at or near those doses that have the most influence on the feature (estimate) of interest.

An example of this strategy is shown in Table 3 which contains calculated values from Equation 9 for the influence of each point on 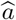 for the data in Fig. The influence values are small and similar for all data points. It is clear from these values that additional data in any region would not affect 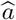 greatly. This is likely due to the relatively substantial amount of data supporting 4 parameter estimates and the good fit of the model. The final (rightmost) data point has the largest influence in the set. As in an ordinary linear regression, its location at the extreme would be expected to have the most leverage on the model slope to the right of the maximum, and consequently on the location of the maximum. Even so, it does not appear worthwhile to gather more data at high doses.

**Table 3.** Influence of individual data values, ordered by dose, on the estimated peak and model fit for the example in Fig. Calculations are defined by Eq 9. Data from envelopes were excluded when calculating these values.

| Data Point | Dose | Influence $\lambda_j$ | Error Variance | Adjusted $R^2$ | AICc* |
| --- | --- | --- | --- | --- | --- |
| 1 | 0 | 0.033 | 0.019 | 0.981 | 1.65 |
| 2 | 10 | -0.004 | 0.020 | 0.980 | 2.31 |
| 3 | 15 | 0.003 | 0.021 | 0.978 | 3.10 |
| 4 | 30 | -0.022 | 0.021 | 0.980 | 3.03 |
| 5 | 40 | -0.002 | 0.022 | 0.974 | 3.46 |
| 6 | 50 | 0.005 | 0.022 | 0.972 | 3.42 |
| 7 | 55 | 0.019 | 0.020 | 0.974 | 2.50 |
| 8 | 60 | 0.012 | 0.021 | 0.974 | 2.91 |
| 9 | 75 | -0.007 | 0.021 | 0.980 | 2.91 |
| 10 | 85 | -0.001 | 0.022 | 0.976 | 3.42 |
| 11 | 100 | 0.000 | 0.017 | 0.983 | 0.98 |
| 12 | 100 | 0.000 | 0.018 | 0.981 | 1.20 |
| 13 | 125 | -0.066 | 0.006 | 0.995 | -14.2 |

\*AICc is the finite sample corrected Akaike Information Criterion.

#### Effects on overall model fit

The set of 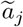 constructed by removing each data point one at a time is a jackknife set [8]. An alternative to calculating point influences on the feature of interest is to calculate model fit summary statistics for each jackknife sample. More data might then be gathered in the region that improves model fit the most. It is not obvious which of many model fit statistics is the best basis for such judgments. Three examples are shown in Table 3, the overall error variance, *R*^2^ adjusted for the number of parameters, and the finite sample corrected Akaike Information Criterion (AICc).

These model fit statistics indicate also a focus on the highest dose point. When it is excluded, the error variance decreases, the *R*^2^ increases, and the AIC indicates improvement in the overall model fit. It is thus somewhat inconsistent with the remaining data. This might be grounds for repeating the dose. Like the influence measures, model fit statistics may suggest gathering new data in regions distant from the feature of interest.

#### New design points

To illustrate the second optimization method, forward looking simulations can be used. We can do this by specifying a new small envelope that represents a hypothetical location for the next design point, simulating data from it, and observing the effect on the feature of interest. Varying the location of the hypothetical design point might suggest an optimal position, in which case real data could be gathered there. For such simulations, the envelope would likely be small and located on the apparent dose response curve. A single data point per simulation with the same weight as real data would be used. This process is illustrated in Fig for a saturation model given by Equation 6. The optimal location for the next design point might be the one that induces the largest variability in the estimand, because this implies it has the most influence on the result.

A key concept is that testing a hypothetical new design point using an envelope does not employ the model to generate data for simulation. It does use the model for fitting. However this process is free of assumptions about true model parameters or the error structure of observations around the model. Hence it is potentially more flexible and robust than ordinary simulations.

### Operating Characteristics

Operating characteristics (OC) are the performance of an algorithm when the truth of nature is known (assumed). A method that performs well under a variety of assumptions is likely to reflect the truth when it is uncertain. Conversely, a method that performs poorly under known conditions is also likely to perform poorly under unknown conditions. The weights for envelope samples used by EnSim are so small compared to those for real data that the algorithm essentially becomes a CRM late in its cycles, known to have good OCs. In the important but narrow class of dose finding questions where a maximum tolerated dose (MTD) is sought, CRM methods are known to perform better than operational designs such as the 3+3.

We have compared the EnSim and 3+3 design OC for a small sample of MTD problems. Space does not permit reporting an extensive series of simulations. However, this comparison is partly beside the point because EnSim is designed to apply outside the MTD context where there are no general tools. Fig shows a typical example of OCs for EnSim compared to a 3+3 design for MTD determination. The true dose response function is shown in blue where the 0.33 quantile is 48. A highly reliable design will stop close to that on average. A very square OC is desirable as it indicates little variation in stopping close to the target. In this example, the median EnSim stopping point was 47 with a true probability of 0.31, based on 100 replicates. The EnSim recommended dose was taken to be the median of the dose distribution for every simulation. The 3+3 algorithm showed a 90% chance of stopping at or below 48 based on 1000 replicates. Hence the 3+3 algorithm is strongly biased on the low side. These results are typical of wider behavior.

These OC comparisons are unfair to a poor design such as the 3+3 for the following reason. Because the 3+3 uses a decision rule based on 1 MTD out of a cohort of 3 does not mean that it is titrating to a probability response of 0.33. Many users of this design behave as if it does, yet the OC shows that it does not. For a given true dose response model, it might be possible to adjust the 3+3 doses, cohort sizes, decision rules, and so on to titrate to 0.33 based on extensive simulations. But different details would be necessary if the true dose response model were assumed to be different. There would be no universal approach as there is with model guided algorithms. So the 3+3 algorithm is defective for selecting anything but a conservative dose with unknown properties, and really can’t be compared to model guided dose escalation.

### Dual Outcomes

Because EnSim is not tied to a single dose response model, it can be adopted to relate dose simultaneously to more than one outcome. It is usually difficult to construct a joint model for multiple outcomes, but easier to model them independently over a common dose basis. Although outcomes may be correlated because, among other reasons, they arise in the same subject, dose finding can be guided using such marginal models. For example, suppose a clinical outcome is best represented by probability of response such as Equation 1, and a simultaneous pharmacological outcome is best represented by a saturating model such as Equation 6. Assume that optimal doses can be characterized for each outcome. We could use the clinical response model to titrate dose conditional on the predicted response from the pharmacologic model being above (or below) a set threshold. Both outcomes would be measured in each subject at every step of the trial, and the models fit independently to their respective data to provide the best estimate of each dose response relationship.

If the primary model predicts a new dose that violates the threshold in the secondary model, the dose will be adjusted accordingly. Such a strategy does not explicitly model the relationship between the two outcomes but allows each to influence the dose estimated. It seems unlikely that a single dose can be optimal for two outcomes. But it is reasonable to assume that a single dose can be optimal for one outcome while simultaneously being acceptable for another. If no such dose can be found, the therapy would have to be judged ineffective. This approach for two outcomes is feasible using EnSim with sufficient data, and can avoid having to construct a joint model for simultaneous outcomes.

## Discussion

Dose finding in its broad form is a challenging methodologic problem. Aside from ethics constraints surrounding the use of a new therapeutic in human subjects, dose finding trials are usually small, and can yield unexpected behavior at high doses of a new therapeutic. Also, the optimum dose being sought almost certainly has different characteristics in various clinical settings or for different classes of therapeutics. Despite these difficulties, two versatile components that yield both efficiency and flexibility for a general solution are a plausible biological model to guide dose escalation in the presence of real data, and statistical simulations to represent imperfect knowledge and biological variability.

EnSim demonstrates the potential value of these elements in many hypothetical examples. The EnSim algorithm can be understood as a generalization of the continual reassessment method. We therefore expect the method to reflect efficiency and unbiased characteristics of model guided dose finding. As a consequence we have not conducted extensive comparisons of EnSim to other dose escalation methods for MTD determination. The limited work on this topic we report in this paper is exactly as expected. Better operating characteristics do not seem to the dominant reason for investigators choosing a design in any case. But EnSim is simple and flexible, making it immediately applicable for problems where there is no standard approach. There is nothing to compare it against for the general class of dose optimization problems.

EnSim is not a Bayesian method strictly speaking. It has more of the spirit of multiple imputation for missing data. However, there is a strong conceptual similarity between assuming a prior probability distribution for unknown model parameters in a true Bayesian framework versus specifying pseudo-data to accomplish the same aim. Fitting pseudo-data with a model explicitly yields parameter estimates with an implied joint probability distribution. For a statistician this may seem an awkward way to reach a “prior” probability distribution, but for clinical investigators it may fit intuition very much better.

Aside from being a method to find the dose optimum in a specific biological question, EnSim contains the framework to improve the experiment design during its conduct. While this is a secondary goal and might be approached in various ways, the creation of a model, a stable fit to real data, and a framework for simulation opens a door to optimizing the design along with the dose. It remains to be seen if this approach can simplify complex design problems. However it does appear perfectly feasible to sharpen up a dose finding design near its conclusion, for example by adding a small number of design points if they appear likely to improve estimation.

In conclusion, EnSim is a new method but familiar in the domain of model guided dose finding. It is simple and flexible, and sure to be operationally superior to empirical designs. It may also be relatively easy for clinical investigators to embrace. For these reasons we believe this method deserves practical application to assess its strengths and weaknesses.

## Figure Legends

**Figure 1.**
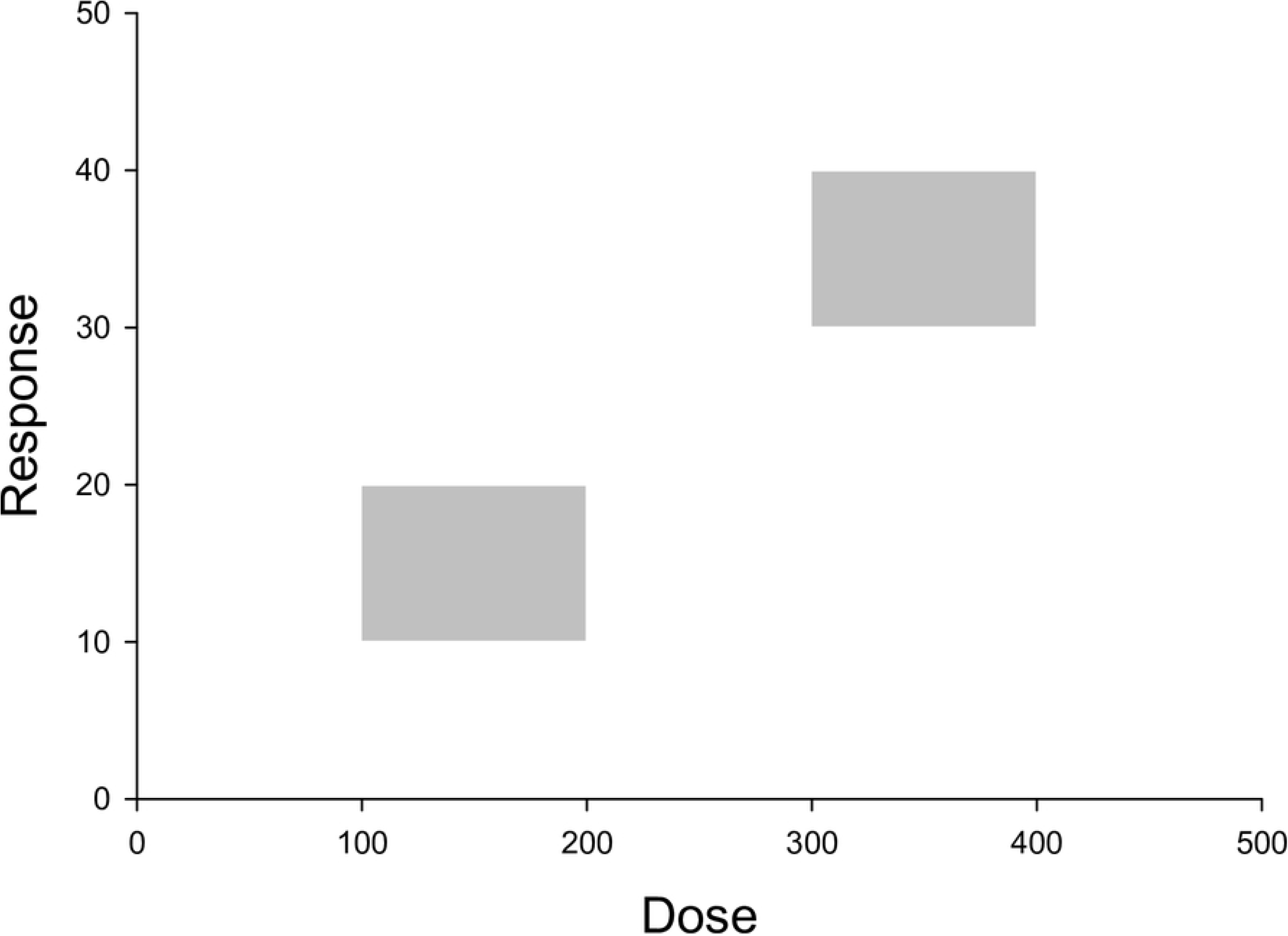
Envelopes representing hypothetical regions of data in a dose-response plane. The dose response model should pass through these regions, absent real data to the contrary. Envelopes such as these represent both what is known and unknown about dose response early in the solution process.

**Figure 2.**
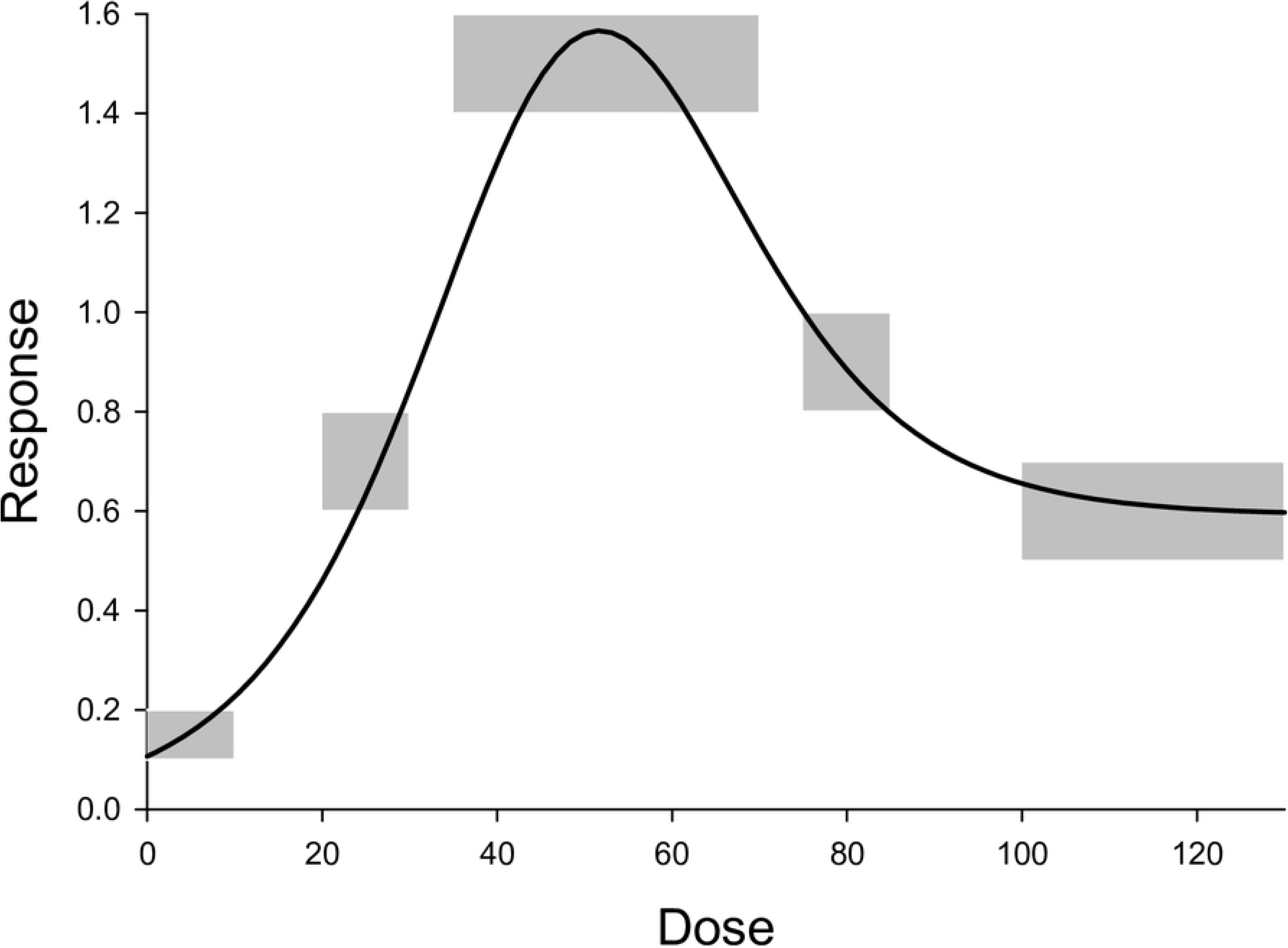
Hypothetical envelopes in a dose response plane and a model consistent with them. The line is a representation of the model given by Eq 2.

**Figure 3.**
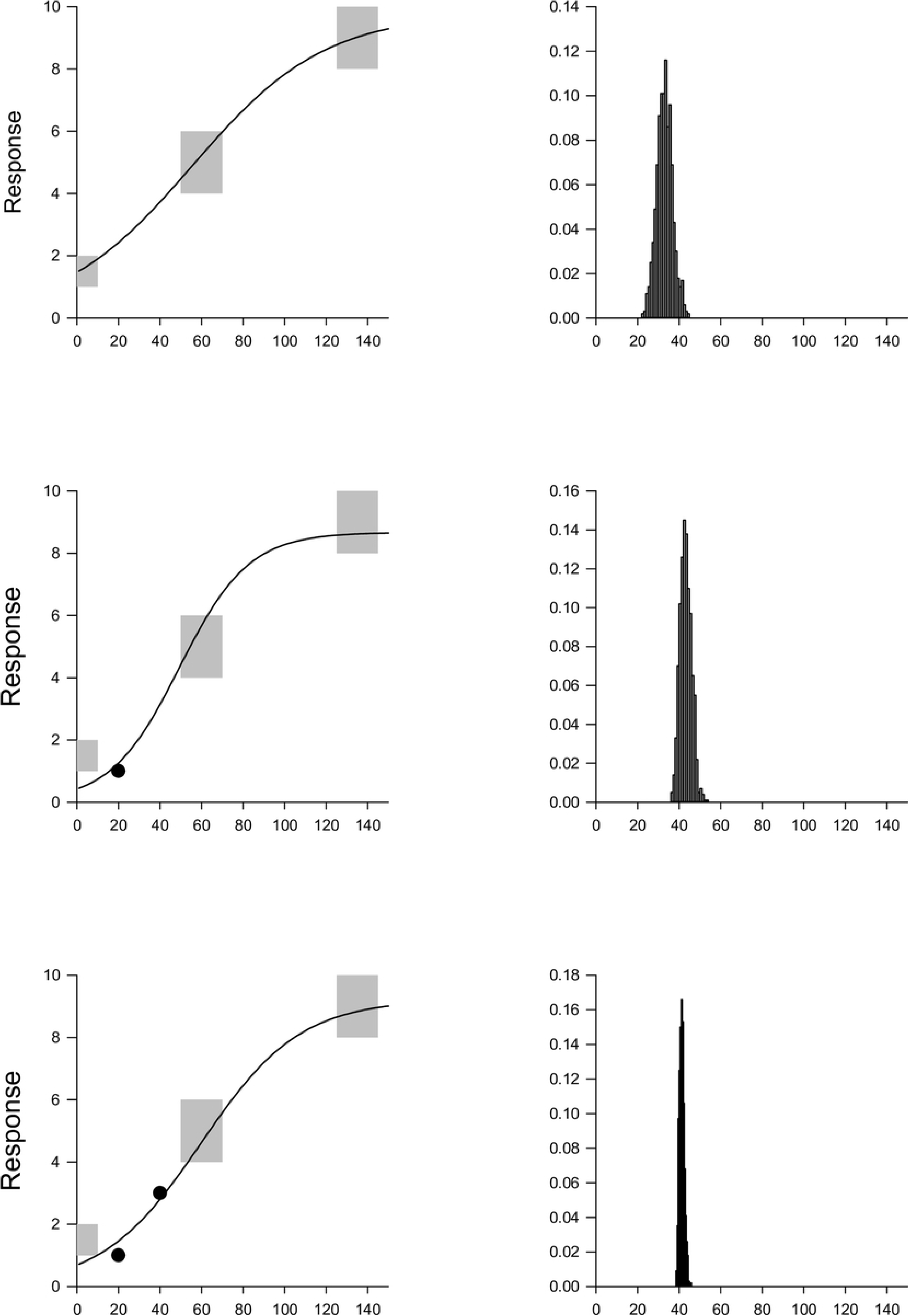
First 3 steps of a hypothetical EnSim dose escalation using the logistic model of Eq 1. The left column shows dose versus response. The right column shows the histogram of estimands from replicates in the simulation. The top row shows Step 1 or initiation based on envelope data only. The histogram of doses associated with response 4.0 is broad. The middle panels show Step 2 and the impact of the first real data point (circle). The histogram is narrowed and shifted to the right, consistent with the fitted model. The bottom panel shows Step 3 and support from two data points, where the estimand histogram narrows. Real data are beginning to dominate the model fit. Subsequent steps are shown in Fig.

**Figure 4.**
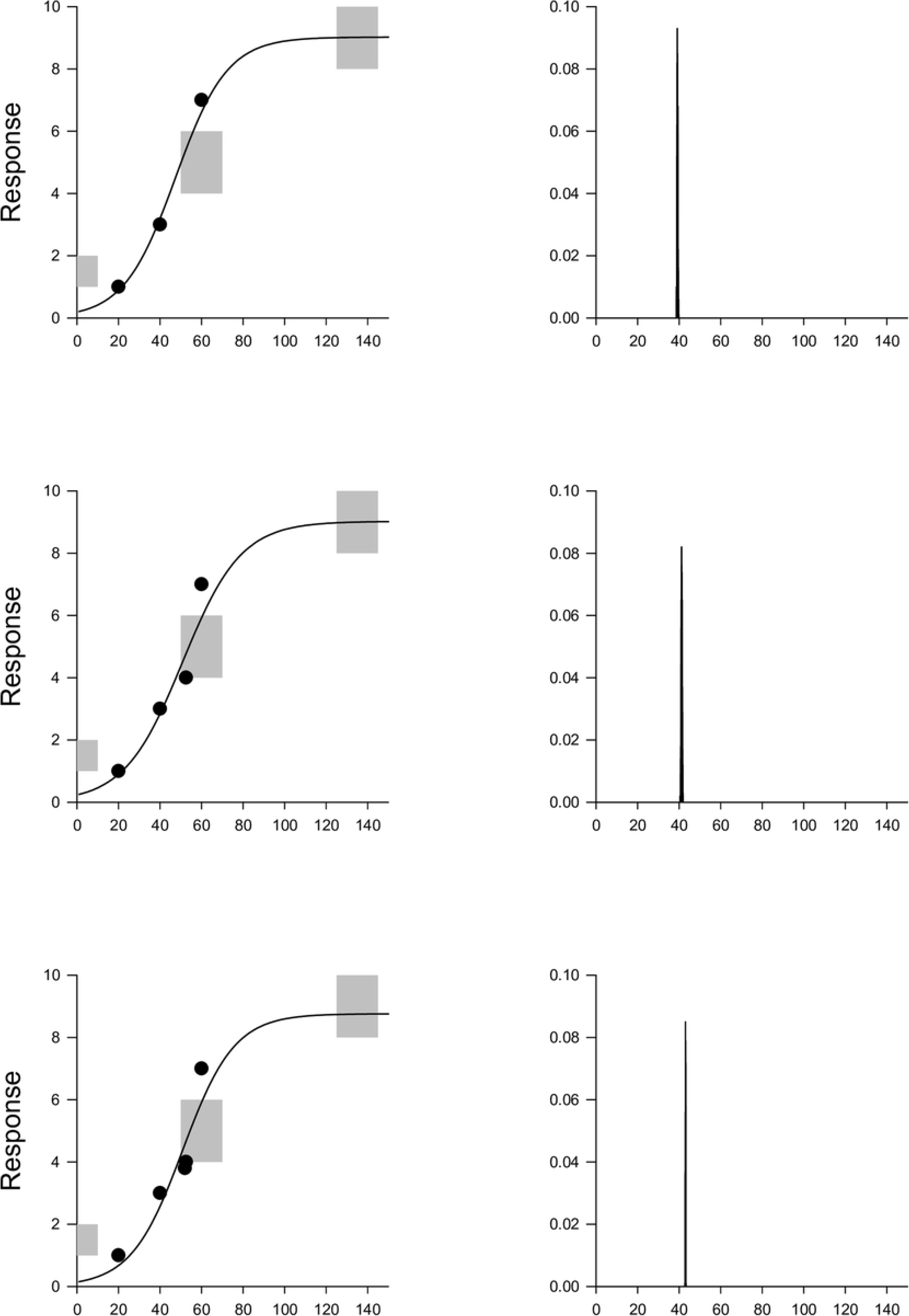
Final 3 steps of the hypothetical EnSim dose escalation from Fig using the logistic model of Eq 1. The influence of simulated data from the envelopes disappears as evidenced by narrowing histograms for predicted dose, and the final model fit is determined almost entirely by real data (circles). Parameter estimates for the model are given in the text.

**Figure 5.**
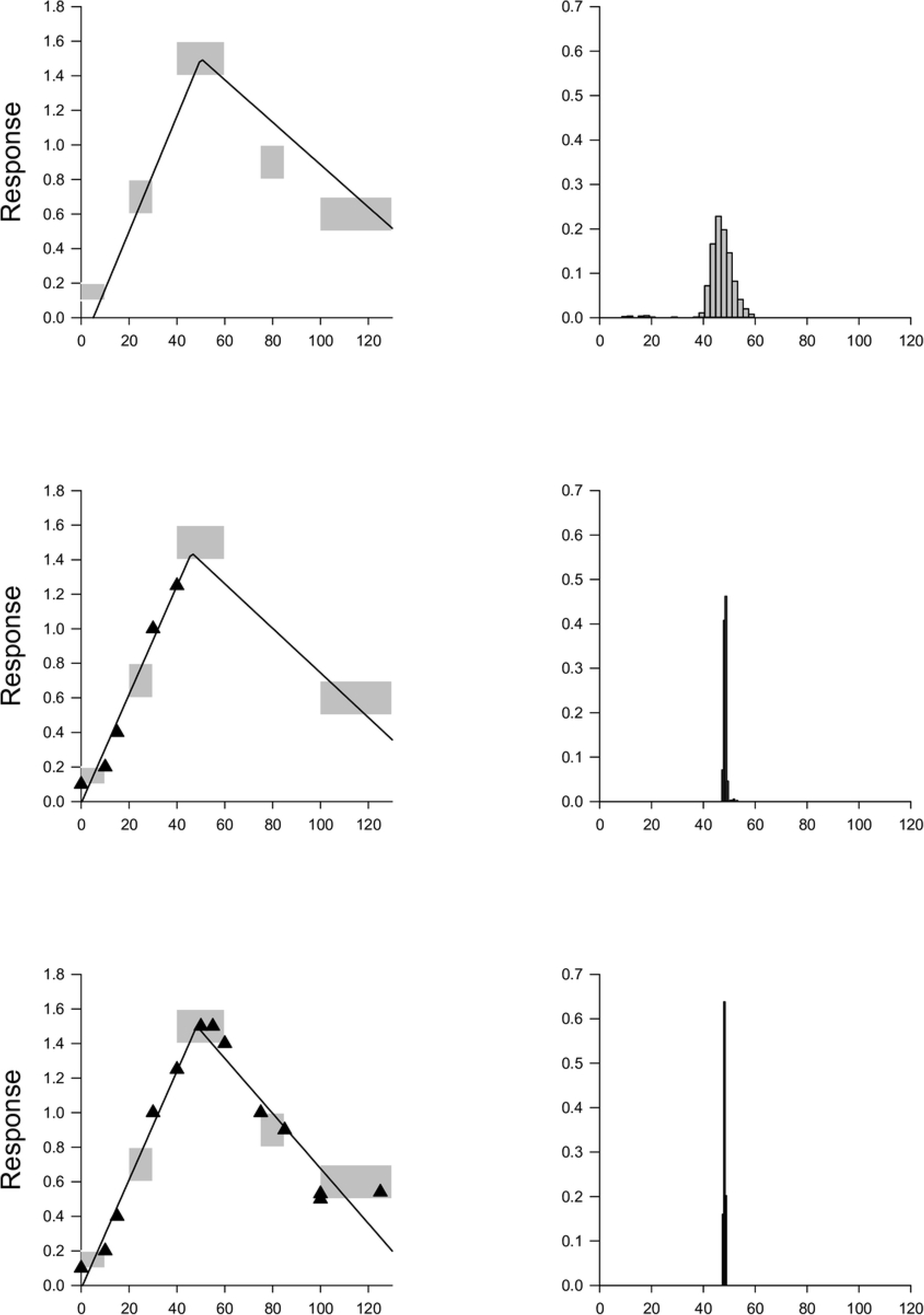
Example of EnSim model fits using Eq 5. The left column shows dose versus response. The right column shows the corresponding histograms of estimated peak values from the simulations. The initial fit (top panel) is based on envelope data only. Data gathered from below the peak are shown in the middle panel (triangles). The full fit of all data is shown in the bottom panel. Estimated parameter values for the model are shown in Table 2.

**Figure 6.**
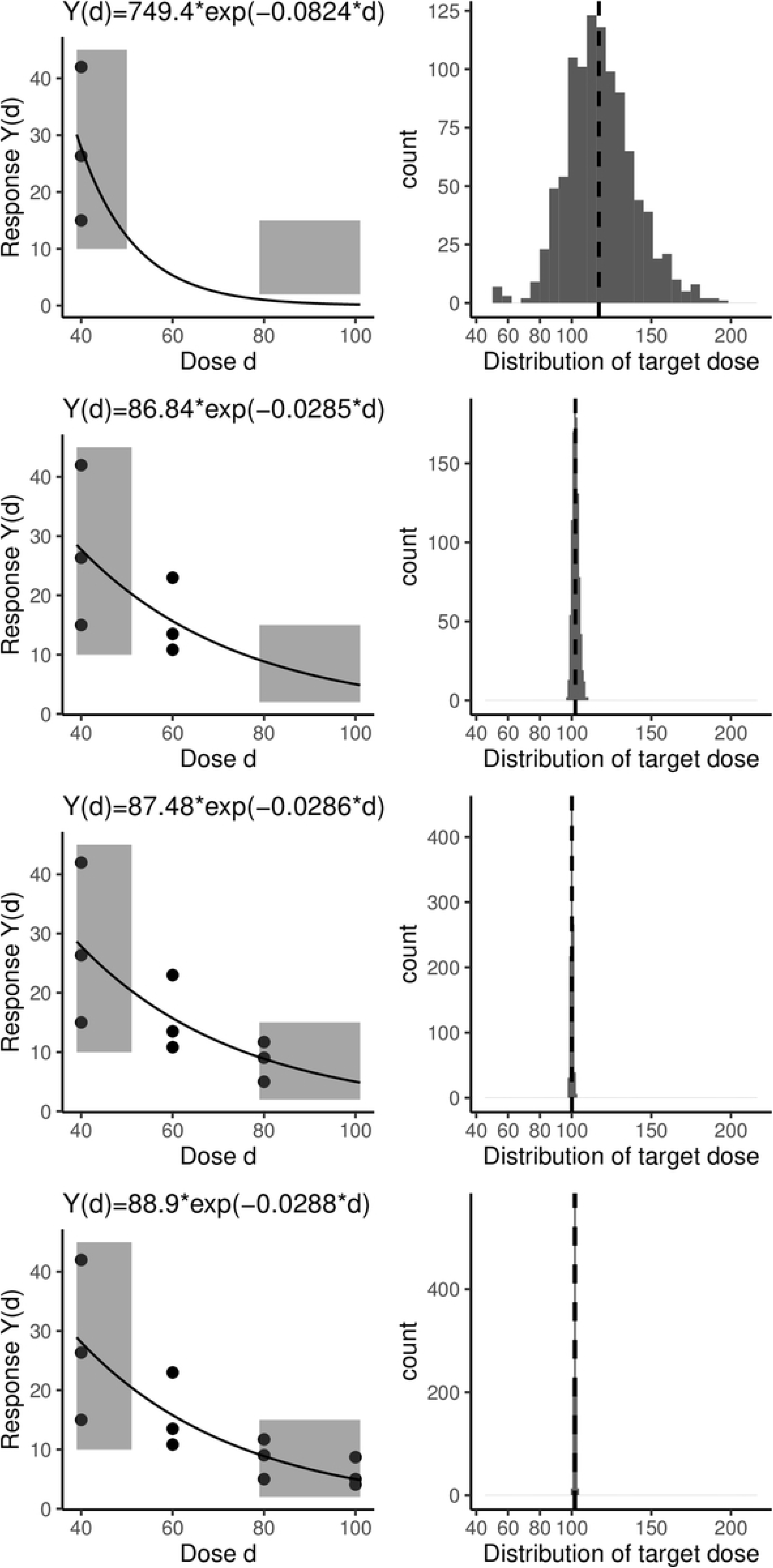
EnSim applied to the trial data from Table 1 in Ivanova and Kim [10] (resampled data from Friedman et al. [9]) as explained in the text. The histograms are based on 1000 simulations. The model recommended dose is indicated by a dashed line in the histogram. The solid dots are the real data and the curves represent the fitted dose-response models.

**Figure 7.**
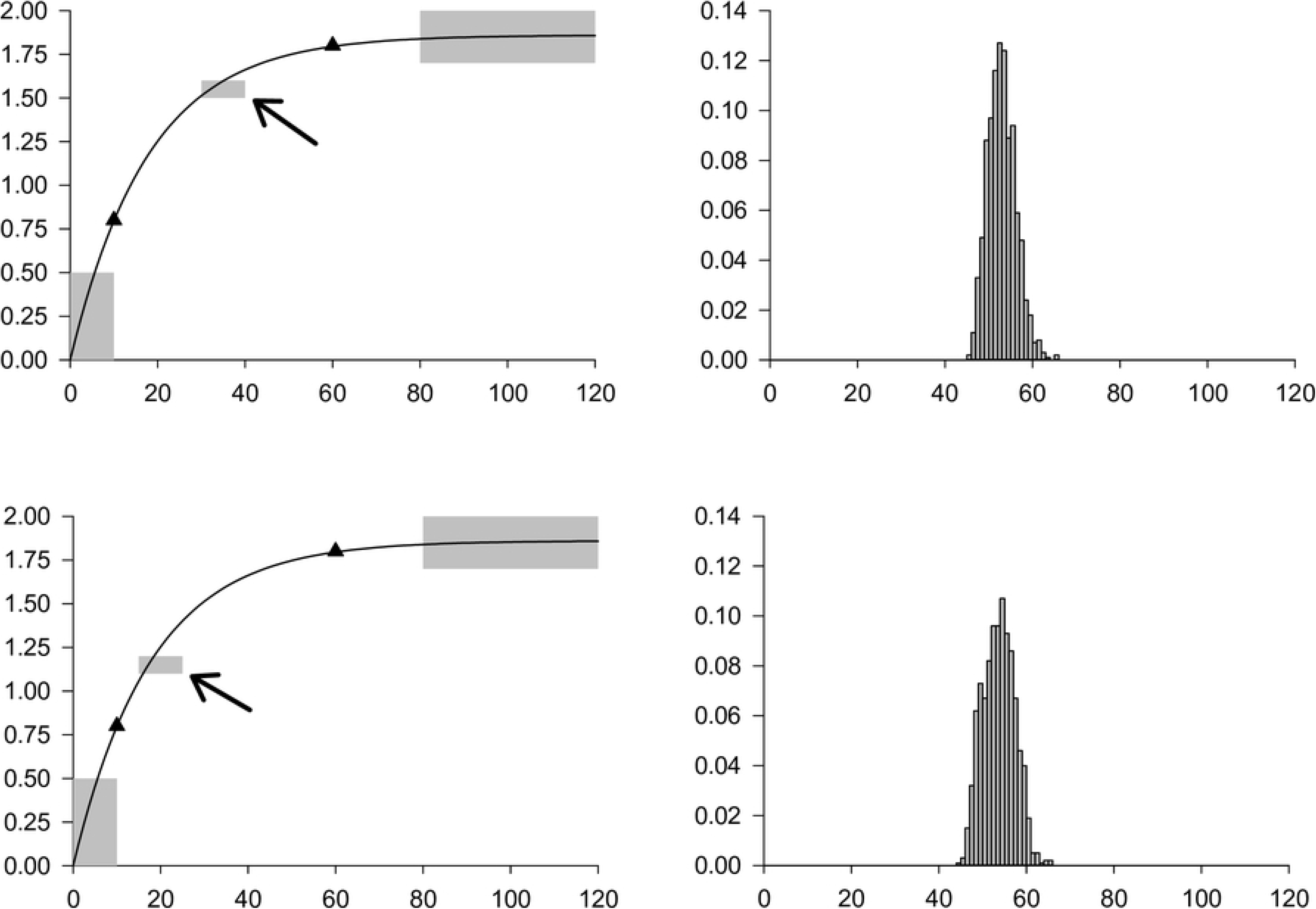
Simulations to determine an optimal location for the next design point in a model given by Eq 6 after two doses have been tested (triangles). The dose yielding a response of 1.75 is sought. A design point close to the optimum (upper left panel) produces less variability in the estimated dose than a design point at lower dose (lower left panel) as indicated by the histograms (right). Consequently the lower dose appears to have more leverage and might be the better choice. Results are based on 1000 replicates of a single data point simulated from the indicated envelope.

**Figure 8.**
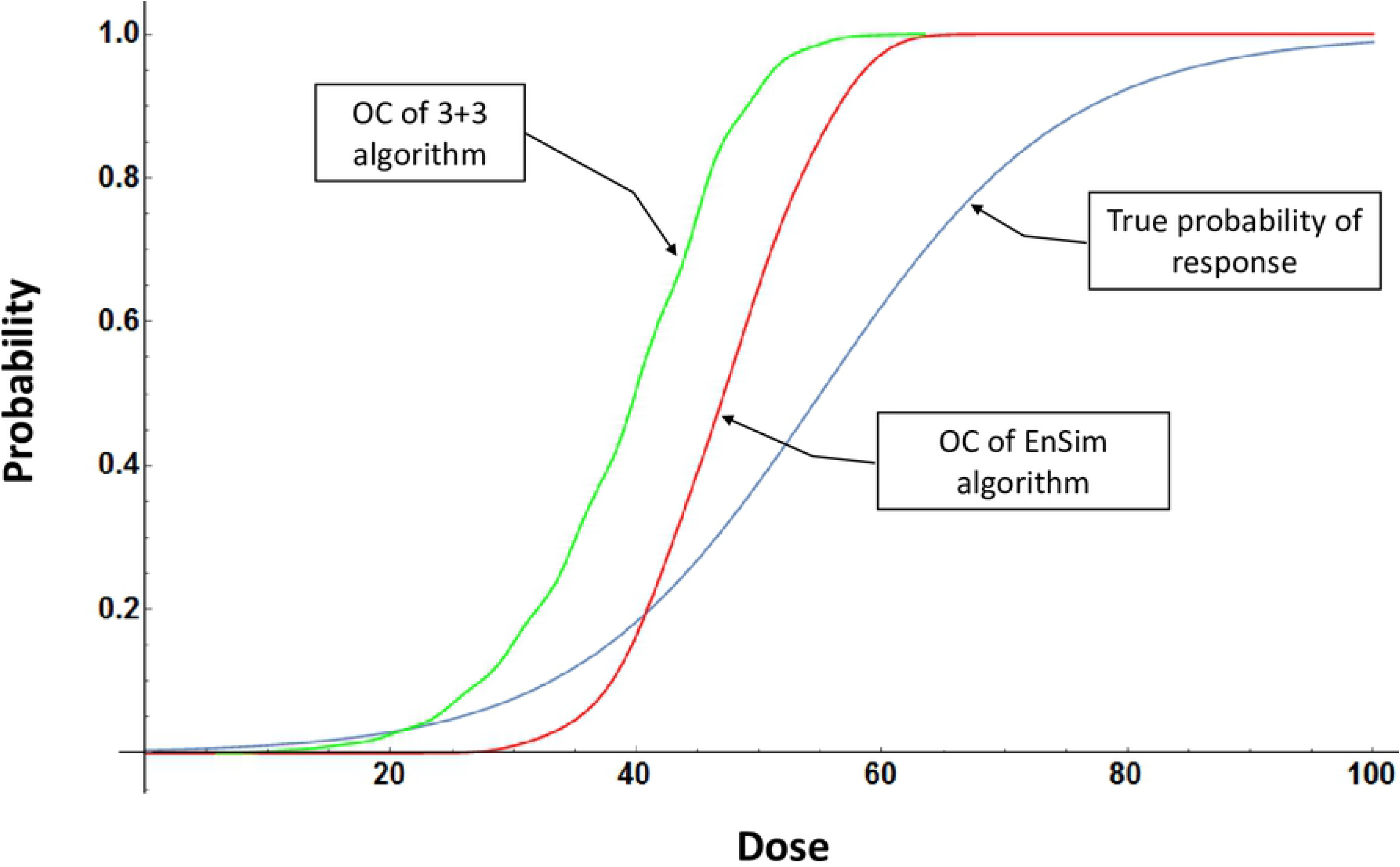
Example operating characteristics of EnSim versus 3+3 algorithm. Results are based on 100 EnSim replicates (red) and 1000 3+3 simulations (green) using the specified true dose response function (blue). The true 0.33 quantile of the dose response function is 48; the median of the EnSim simulations is 47; 90% of the 3+3 simulations terminate at a dose lower than 48.

## Supporting Files

1. R code for Example 2 and Fig 6.
2. Mathematica code for general EnSim model fitting and Fig 1-5.
3. Mathematica code for operating characteristics and Fig 8.

